# OPERA-LG: Efficient and exact scaffolding of large, repeat-rich eukaryotic genomes with performance guarantees

**DOI:** 10.1101/020230

**Authors:** Song Gao, Denis Bertrand, Burton KH Chia, Niranjan Nagarajan

**Affiliations:** Computational and Systems Biology, Genome Institute of Singapore, Singapore 138672, Singapore; NUS Graduate School for Integrative Sciences and Engineering, Singapore 117456, Singapore; South Australian Health & Medical Research Institute, North Terrace Adelaide 5000 SA, Australia

**Author notes:** Contributed equally (Song Gao, Denis Bertrand). Correspondence: Niranjan Nagarajan.

## Abstract

The assembly of large, repeat-rich eukaryotic genomes continues to represent a significant challenge in genomics. While long-read technologies have made the high-quality assembly of small, microbial genomes increasingly feasible, data generation can be prohibitively expensive for larger genomes. Advances in assembly algorithms are thus essential to exploit the characteristics of short and long-read sequencing technologies to consistently and reliably provide high-quality assemblies in a cost-efficient manner. OPERA-LG is a scalable, exact algorithm for the scaffold assembly of large, repeat-rich genomes, with consistent improvement over state-of-the-art programs for scaffold correctness and contiguity. It provides a rigorous framework for scaffolding of repetitive sequences and a systematic approach for combining data from different second-generation (Illumina, Ion Torrent) and third-generation (PacBio, ONT) sequencing technologies. OPERA-LG efficiently scaffolds large genomes with provable scaffold properties, providing an avenue for systematic augmentation and improvement of 1000s of existing draft eukaryotic genome assemblies.

## Background

The field of sequence assembly has witnessed a significant amount of mathematical and algorithmic study of the problem [1–4]. Yet many assembly programs in use do not have a clear objective function that they optimize, relying on heuristics and/or manually-tuned parameters to piece genomes together to variable degrees of success [5–7]. As there are a wide array of heuristics and parameter choices to try, the right combination that works well across a range of datasets may not always be apparent and new assembly tools run the risk of being tuned for the datasets on which they are benchmarked. Recent assembly competitions such as GAGE [8], Assemblathon [9], Assemblathon2 [10], and a recent scaffolder benchmark [11] have thus played an important role in galvanizing the community and in highlighting the drawbacks of existing tools.

The prevalence of heuristic choices in assembly owes its origins partly to several well-known early results regarding its computational complexity [1, 2] (and further confirmed by recent studies [3, 4]) which suggest that most formal definitions of various assembly problems (such as contiging and scaffolding) are computationally intractable (NP-hard). Notably though, most complexity results have been limited to worst-case analysis and relatively little has been said about average-case or parametric complexity of various assembly problems [1, 4, 12]. For example, while the problem of constructing contigs from read data (typically formulated as a path-finding problem) has been shown to be NP-hard in terms of worst-case complexity [3, 4], in practice, the problem is usually under-constrained in the absence of ultra-long reads, and trivially-computable, fragmented contig assemblies are the best we can do [4, 12]. The use of paired-end and mate-pair reads to scaffold contigs thus plays a vital role in assembly projects to significantly boost assembly quality [13–16]. While worst-case analysis for the scaffolding problem also suggests that it could be computationally expensive to solve exactly, surprisingly, it is possible to design exact algorithms that require runtime polynomial in the size of the scaffold graph [13]. These algorithms guarantee a scaffold assembly that minimizes discordance with the input data and thus provide an internal “quality-control” for their results. As shown in Gao *et al*. [13] using experiments on small genomes, an exact algorithm for scaffolding simultaneously leads to more accurate as well as contiguous assemblies.

As the sequencing process is stochastic, in principle, probabilistic models and objective functions provide a natural approach for assembly and scaffolding [16–20]. Methods such as Genovo [19] and SOPRA [16] are based on such models and provide an alternative approach that is particularly powerful when sequencing coverage is low and read information uncertain. When sequencing coverage is high (>20X, as is often the case for short-read genome assembly projects) noise from random chimeric reads is easier to filter. Correspondingly, scaffold assembly is frequently formulated as a combinatorial graph problem and this is the approach followed in this study.

Based on the scaffold assembly formulation from Huson *et al*. [21], this study proposes extensions to an exact algorithm [13] that make it feasible for scaffolding large, repeat-rich genomes in a time and memory-efficient manner [22, 23]. The new approach, termed OPERA-LG, was extensively evaluated against state-of-the-art scaffolders (SSPACE [14], SOPRA [16] and BESST [24]) and assembly pipelines (SOAPdenovo [25] and ALLPATHS-LG [26]) on simulated and real datasets. In the presence of multiple and large fragment (>4 kbp) mate-pair libraries, OPERA-LG was seen to provide several-fold increase in contiguity metrics and/or reduction in scaffold errors. Improvements with a single short-fragment mate-pair library were limited, but OPERA-LG provided consistently good assemblies in these cases as well (among the top 3 scaffolders).

OPERA-LG incorporates several features useful for producing high quality draft assemblies for large, repeat-rich genomes. These include the ability to simultaneously use data from multiple libraries [27] as needed in large assembly projects, an improved edge-length estimation algorithm and an exact extension for scaffolding repetitive sequences that typically confound assembly tools. OPERA-LG’s ability to be sequencing platform independent was evaluated using PacBio and ONT data and compared to scaffolders such as SSPACE-LongRead [28] and LINKS [29].

## Results

### Overview

The algorithmic core of OPERA-LG is adapted from the approach described in Gao *et al*. [13] (summarized in **Supplementary Figure 1**) and is based on (i) a memoized-search to find a scaffold that minimizes the number of discordant read-derived-links connecting contigs (**Supplementary Figure 1a**), (ii) a graph contraction technique that allows for localizing the search for an optimal scaffold without losing the guarantee of a globally optimal scaffold (**Supplementary Figure 1b**) and (iii) a quadratic programming formulation to compute gap sizes that best match mate-pair derived distance constraints [27] (**Supplementary Figure 1c**). To enable it to produce long and accurate scaffolds for large, repeat-rich genomes, OPERA-LG incorporates several novel features and improvements including (a) Optimized data-structures to improve its scalability (b) Refined edge-length estimation and the ability to simultaneously use multiple-libraries to improve scaffolding accuracy and (c) Extensions that allow for the scaffolding of repeat sequences. Algorithmic details for each of these features and improvements in OPERA-LG can be found in the **Methods** section.

**Figure 1:**
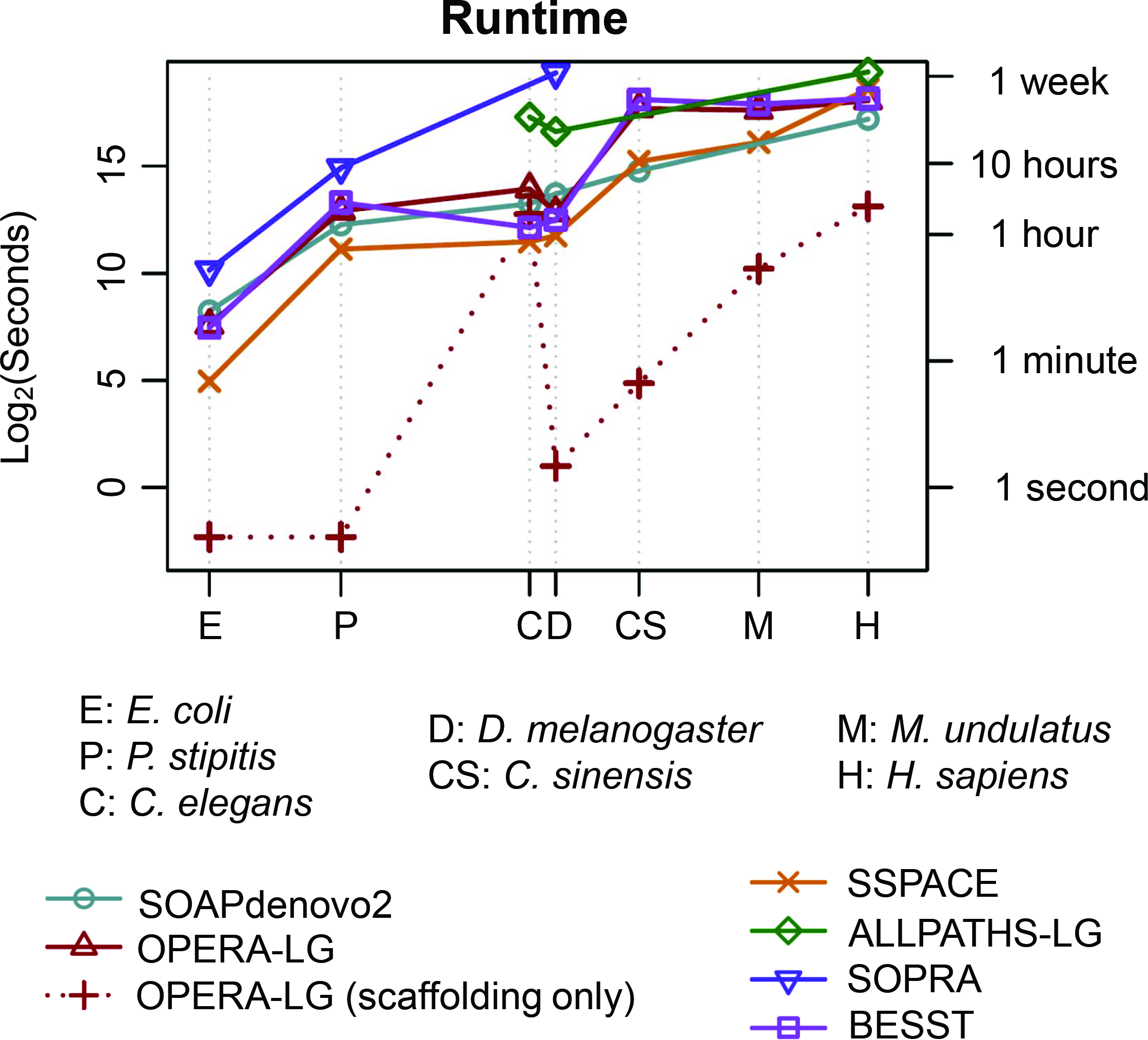
**Runtime as a function of genome size**. Note that both the x and y-axes are log-scaled and the results for various genomes are indicated at the corresponding genome size. For all standalone scaffolders, runtimes include the mapping stage. For SOAPdenovo2 and ALLPATHS-LG we report the runtime of the full assembly pipeline (including contig assembly). “OPERA-LG (scaffolding only)” shows the runtime of the scaffolding algorithm in OPERA-LG, excluding preprocessing, to highlight that it can take a fraction of the overall runtime and is influenced more by the repeat complexity of the genome than the size of the genome. Due to its library-size restrictions, ALLPATHS-LG could be run on only a few of the datasets shown here. SOPRA runs did not complete after 10 days for *C. elegans* and the 3 largest genomes.

## Scalability and Multi-library Scaffolding

### Scalability

Runtime and memory optimizations in OPERA-LG (see **Methods**) are key to its scalability as shown in **Table 1**. In particular, while the method in Gao *et al*. [13] was unable to scaffold the full *D. melanogaster* dataset due to excessive memory usage, OPERA-LG takes a few seconds and a few hundred megabytes of memory (largely for storing read mapping information). For genomes where the previous method was feasible, OPERA-LG is typically >10 times faster and requires significantly less memory (roughly 1/2 to 1/20^th^ of the requirements for Gao *et al*.). Across datasets, OPERA-LG’s runtime was found to be comparable to SOAPdenovo2 and the scaffolders BESST and SSPACE, while requiring less runtime than SOPRA (runs on *C. elegans* and the 3 largest genomes were stopped after 10 days) and ALLPATHS-LG (**Figure 1**). In addition, for small genomes OPERA-LG’s runtime is dominated by the preprocessing step, while runtime for the core of the algorithm (see **Figure 1**, “scaffolding only”) may not necessarily increase in proportion to the genome size (i.e. may be determined by intrinsic features of the genome such as repeat lengths and distribution). Overall, OPERA-LG’s runtime was less than 1 day (on a single processor) using <60 GB of memory for all the datasets tested here (including the human genome), establishing its feasibility for scaffolding large genomes and retaining the potential for further improvement with parallelization.

### Edge Length Estimation and Multi-library Analysis

OPERA-LG was redesigned to simultaneously use data from multiple “jumping” libraries for scaffolding, a process that can be critical for improving assembly contiguity and correctness in large genome assembly projects. These improvements were directly evaluated to establish their utility in OPERA-LG. Firstly, scaffold edge length estimates from OPERA-LG were compared to the known true edge lengths for the synthetic datasets and were found to be in excellent agreement overall (**Figure 2b** and **Figure 2d**). In addition, OPERA-LG’s estimates were found to be more accurate than a commonly used naïve estimation procedure (**Figure 2a** and **Figure 2c**), which was found to consistently under-estimate edge lengths for longer edges, though the bias observed here was not as severe as observed previously [30]. Secondly, OPERA-LG’s ability to handle multiple libraries simultaneously was found to provide a clear benefit over the commonly employed hierarchical approach (**Figure 2e** and **Figure 2f**). The *simultaneous* approach not only led to fewer assembly errors (**Figure 2f**) but also provided improved assembly contiguity (measured by corrected N50; see **Methods**) as a by-product (**Figure 2e**). Note that the results for *C. elegans* may be more indicative of the performance boost that can be expected as the *E. coli* and *D. melanogaster* datasets had high-quality assemblies to begin with.

**Figure 2:**
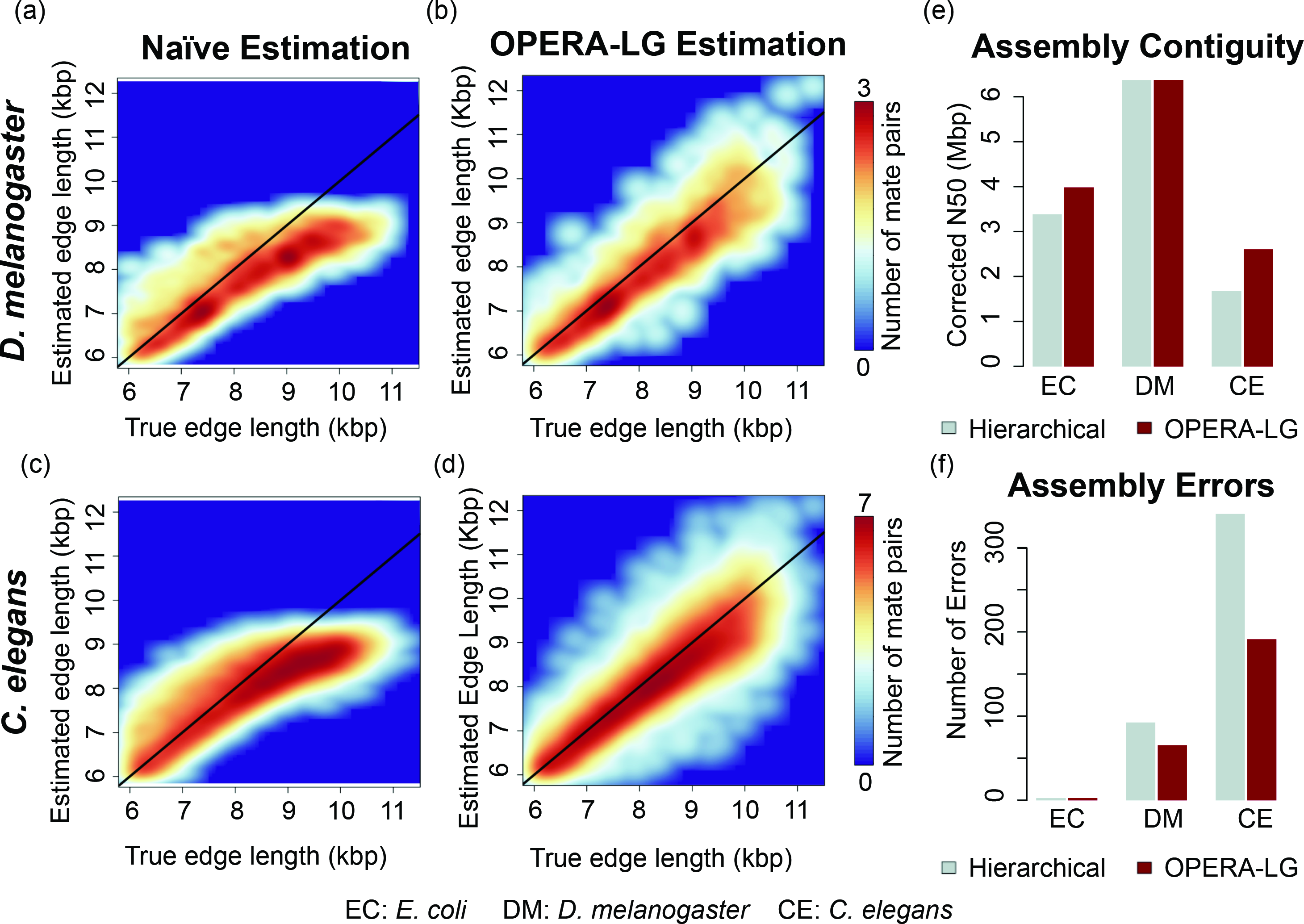
**Improvements in multi-library scaffolding**. Subfigures (a)-(d) show the improved correlation between empirical estimates and true edge lengths when using the procedure in OPERA-LG in comparison to the “naïve estimation” that is commonly used (results reported are for the 10 kbp libraries). Subfigures (e)-(f) depict the improvements in corrected assembly N50 and reduction in corresponding assembly errors when using the multi-library scaffolding implemented in OPERA-LG in comparison to a commonly used hierarchical scaffolding approach that considers libraries independently in order of their insert size.

## Improvements in Assembly Contiguity and Correctness

### Benchmarking with Synthetic Datasets

To evaluate the performance of OPERA-LG, it was first benchmarked on several synthetic datasets as these provide the critical flexibility to vary parameters and assess their effect on the method (**Table 2**). The synthetic datasets contain multiple mate-pair libraries as well as large fragment libraries, representing a typical scenario for the assembly of large genomes where such information is critical (**Table 2**). OPERA-LG was assessed at two levels, a) for its **scaffold quality** by comparing against a recently published method (BESST) and the top three best performing scaffolders from a recent benchmarking paper [11], i.e. SOAPdenovo2’s scaffolding module (S2), SSPACE (SS) and SOPRA, evaluated with a common set of input contigs from SOAPdenovo and b) for **overall assembly quality** (contiging and scaffolding), using SOAPdenovo as a contig assembler and OPERA-LG as a scaffolder (OP), when compared to ALLPATHS-LG (AP) and SOAPdenovo2 (S2) as representatives of state-of-the-art assembly pipelines (i.e. using their contiging and scaffolding modules; see **Supplementary Table 1** for contig and scaffold statistics).

**Table 1:**
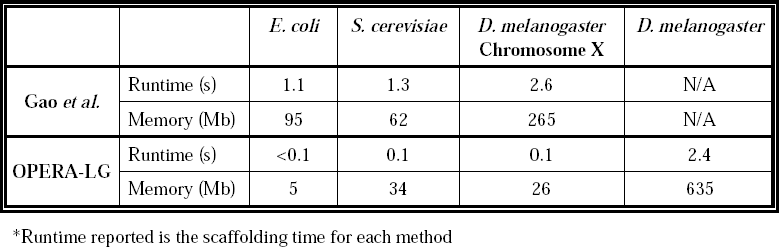
Scalability comparison between the exact method in Gao *et al*. and OPERA-LG.

At the scaffold level, while the corrected N50 for SSPACE, SOAPdenovo2, SOPRA and BESST were typically comparable (**Supplementary Table 2**), OPERA-LG produced assemblies that were significantly more contiguous, regardless of the genome being assembled (5-10× improvement in corrected N50; **Supplementary Table 2** and **Figure 3a**). In addition, scaffolds produced by OPERA-LG contained fewer errors in general (**Supplementary Table 2** and **Figure 3b**), though on the human genome BESST was observed to have fewer indel and relocation errors. Manual inspection of scaffolding errors from OPERA-LG indicate that they were often due to local ordering errors in regions of the scaffold graph that were not sufficiently constrained by scaffold edges (relocations), and gap size estimation errors due to lower read coverage (scaffold indel errors; see **Methods**). In particular, OPERA-LG had few translocation errors, where distant regions of the genome were incorrectly brought together. Results from others scaffolders show that they lead to a large number of translocation as well as inversion (where the orientation of contigs is incorrectly determined) errors compared to OPERA-LG (**Supplementary Table 2** and **Figure 3b**). As such errors can significantly impact downstream analysis this represents an advantage for scaffolds produced by OPERA-LG. It also suggests that the global optimization employed in OPERA-LG may be more effective in eliminating such errors. Overall, OPERA-LG had ≤1 inversion or translocation error for the *D. melanogaster* and *C. elegans* assemblies and less than a 100 such errors for the *H. sapiens* genome.

**Figure 3:**
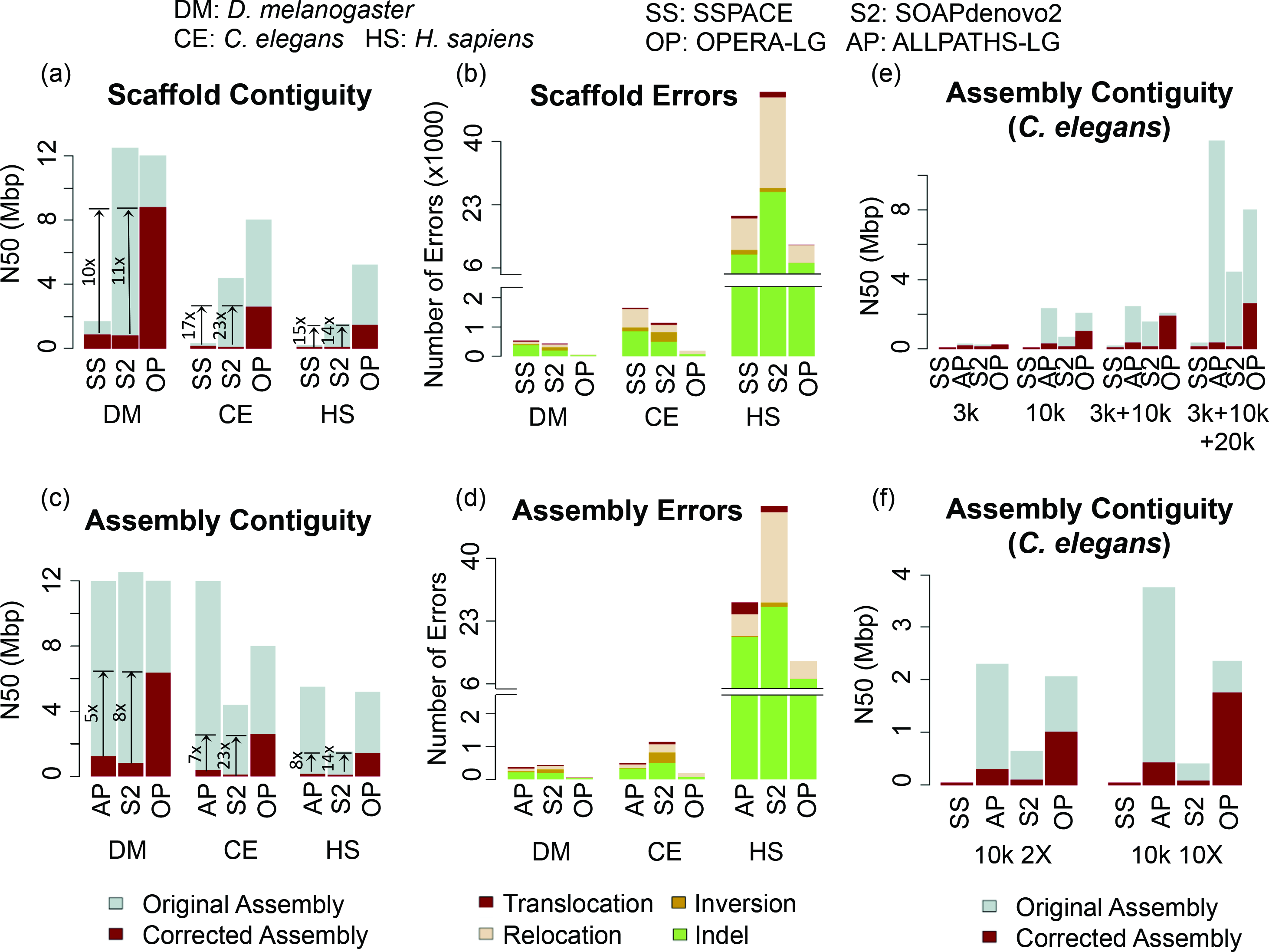
**Boosting assembly contiguity and correctness with OPERA-LG**. Subfigures (a)-(b) show scaffold contiguity and correctness for various scaffolders on different genomes starting from common sets of contigs (generated by SOAPdenovo). The final assemblies were only corrected for scaffold errors. Results for BESST and SOPRA were comparable to SSPACE and are reported in **Supplementary Table 2**. Subfigures (c)-(d) show corresponding overall assembly metrics for various assembly pipelines (that were provided all read libraries as input). The final assemblies were corrected for both contig and scaffold errors to allow for a fair comparison. Arrows highlight fold improvement in corrected N50 using OPERA-LG. ALLPATHS-LG analysis on the human dataset had an abnormal exit after >10 days of runtime but appears to have produced a valid assembly. Subfigures (e)-(f) depict overall assembly contiguity as a function of mate-pair libraries and sequencing depth, provided as input. Results shown are for the *C. elegans* dataset and are qualitatively similar for other datasets as well (data not shown).

**Table 2:**
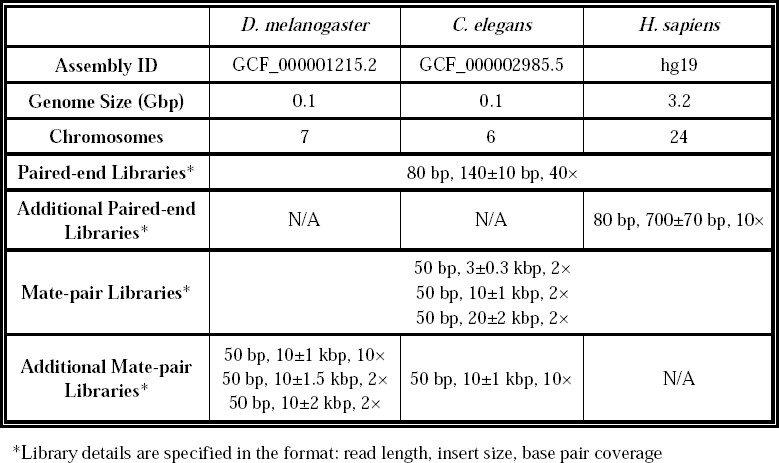
Statistics for synthetic datasets.

For the assembly level comparisons, while ALLPATHS-LG and SOAPdenovo2 had comparable or larger original N50s, corrected assembly N50s were significantly lower compared to OPERA-LG scaffolds based on SOAPdenovo contigs (**Figure 3c**). This was despite the fact that SOAPdenovo contigs were typically more fragmented and thus provided a challenging starting point for the scaffolder. A likely explanation for the improvements seen is the generation of fewer assembly errors compared to ALLPATHS-LG and SOAPdenovo2 (**Figure 3d** and **Supplementary Table 3**). Similar results were seen in experiments with longer reads (**Supplementary Table 4**), indicating that the results are invariant to read length.

OPERA-LG’s performance as a function of the information provided to it was assessed by further studying results for various combinations of libraries as input. Overall, as expected, OPERA-LG was seen to introduce fewer errors when more mate-pair libraries were provided (though this was not necessarily the trend for other methods; **Supplementary Figure 2a**). For example, incorporation of a 3 kbp mate-pair library, in addition to a 10 kbp library, was seen to consistently help eliminate local scaffolding errors in under-constrained regions of the scaffold graph (see **Methods**; **Supplementary Figure 2a**). Correspondingly, both original and corrected N50s improved as OPERA-LG was provided with more libraries (**Figure 3e**). A similar trend was seen with increasing sequencing depth, with OPERA-LG reporting fewer assembly errors (**Supplementary Figure 2b**) and providing consistent improvement in corrected N50 (**Figure 3f)** with higher read coverage.

Finally, the robustness of OPERA-LG to the quality of the sequencing library (measured by standard deviation in library size, with lower values implying higher quality) was assessed. Within a reasonable range of quality, OPERA-LG (and most assemblers) produce very similar assemblies in terms of assembly contiguity (**Supplementary Figure 3a**). However, degrading library quality led to an increase in the number of assembly errors (**Supplementary Figure 3b**). Despite this, OPERA-LG was observed to be more robust to library quality (with fewer errors and better N50 in the worst case than the best-case scenario for other methods; **Supplementary Figure 3b**), confirming its utility for analyzing low quality input data.

**Comparisons on real datasets**. Evaluation of scaffolding performance on real datasets can be influenced by the lack of gold-standard references or limited availability of data. Based on evaluation on six sequenced datasets, OPERA-LG was observed to perform consistently well and provide significant scaffold improvements when multiple and large-fragment libraries were provided as input. For example, for the extensively sequenced parrot genome datasets (*M. undulatus*, estimated genome size = 1.2 Gbp, with 2 kbp, 5 kbp, 10 kbp, 20 kbp and 40 kbp mate-pair libraries), which were assembled as part of the Assemblathon2 competition [10], the corrected N50 obtained using OPERA-LG (with SOAPdenovo contigs) was 2.2× the best reported Assemblathon2 assembly (ALLPATHS-LG; **Figure 4a**). This was despite the original N50 for OPERA-LG being slightly smaller than that for other Assemblathon2 programs, such as ALLPATHS-LG. Notably, OPERA-LG’s results were based on more conservative contigs from SOAPdenovo and using default parameters, while Assemblathon2 results were from the best submissions by their respective teams. In comparison to other standalone scaffolders (that shared the same contig set), OPERA-LG’s corrected N50 was 5.8× that of BESST, and 1.5× that of SSPACE (**Figure 4a**). SOPRA could not be evaluated as it did not complete its analysis even after 10 days of runtime.

**Figure 4:**
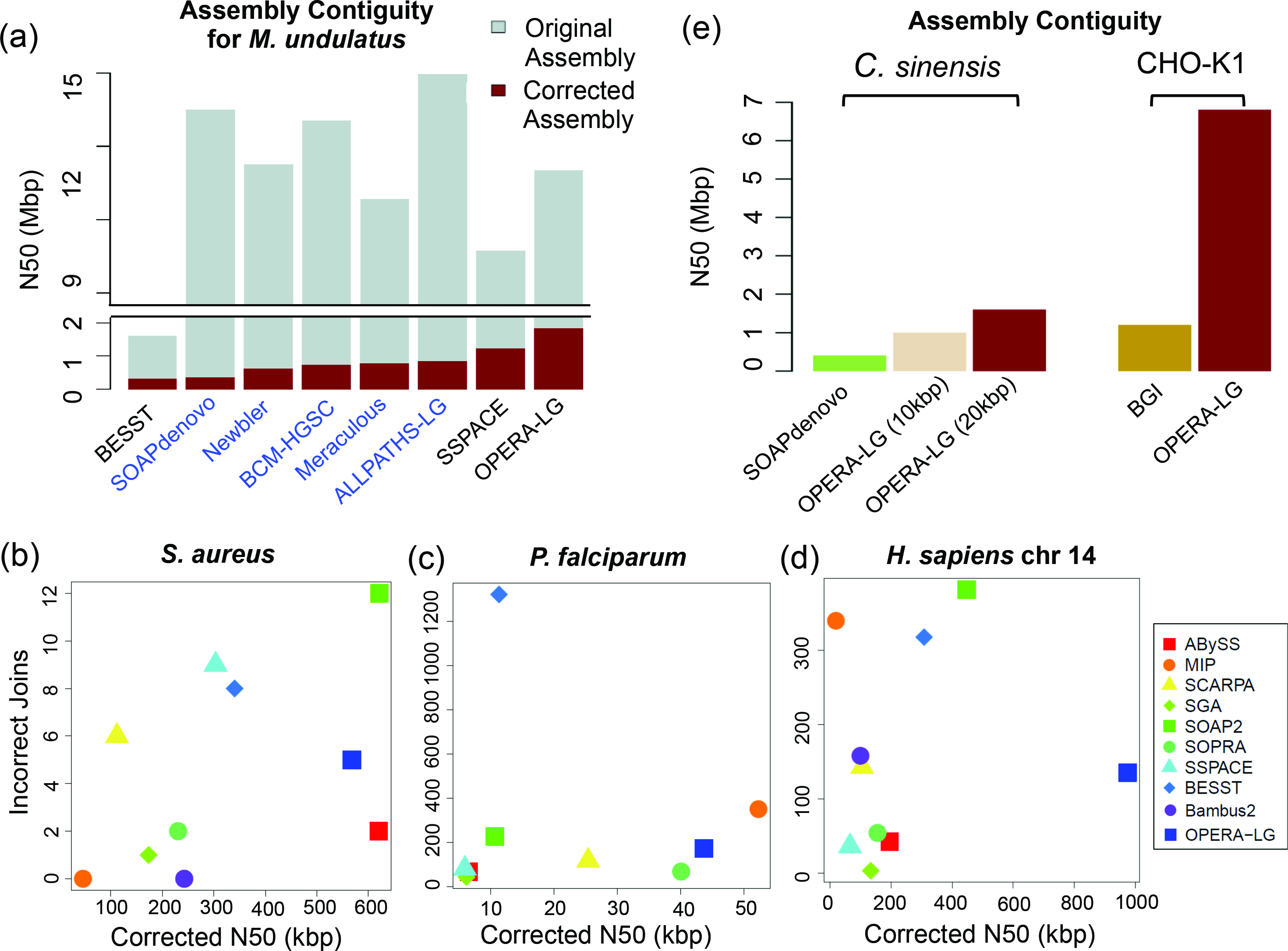
**Assembly improvements on sequenced genomes using OPERA-LG**. (a) Results for the *M. undulatus* genome, comparing OPERA-LG against the best assemblers in the ASSEMBLATHON2 competition (in blue) and standalone scaffolders (in black; SOPRA did not complete after 10 days of runtime). Note that while OPERA-LG’s original N50 is similar to other methods, its corrected N50 is >1.5× that of others. Scatter plot of incorrect joins and corrected N50 for (b) *S. aureus*, (c) *P. falciparum* and (d) *H. sapiens* chromosome 14. OPERA-LG is consistently among the top 3 methods in terms of corrected N50 in this evaluation. (e) Assembly augmentation using OPERA-LG for the *C. sinenis* and CHO-K1 genomes significantly boosts assembly contiguity.

Similar results were obtained for the assembly of a sweet orange genome (*C. sinensis*, genome size = 367 Mbp, with 2 kbp, 10 kbp and 20 kbp mate-pair libraries) [31], where OPERA-LG’s corrected N50 was 2.5× that of the best performing alternative (SOAPdenovo2, 178 vs 68 kbp) and with less than a third of its scaffolding errors (1,492 vs 5,455; **Supplementary Table 5**). SOPRA was unable to run to completion on this dataset after 10 days, while BESST and SSPACE had corrected N50s <60 kbp. In the case of a well-assembled yeast genome (*P. stipitis*, genome size = 15.4 Mbp, 3 kbp mate-pair library), SOPRA ran to completion and was similar to OPERA-LG in terms of producing few scaffold errors (8 vs 1; **Supplementary Table 5**). On the other hand, SOAPdenovo2’s scaffolds were comparable to OPERA-LG in terms of corrected N50 (299 kbp vs 320 kbp) but with many more scaffold errors (61 vs 1; **Supplementary Table 5**). OPERA-LG’s scaffolds improved over SSPACE, BESST and SOPRA in terms of corrected N50 (1.4-3×) and had fewer errors than SSPACE and BESST (**Supplementary Table 5**). In comparisons on the *M. undulatus*, *C. sinensis* and *P. stipitis* genomes, the contigs provided as inputs to the scaffolders were not corrected for assembly errors, and thus these evaluations should better reflect scaffolder performance in real genome assembly projects. In all three cases, OPERA-LG improved corrected N50 significantly, while having fewer scaffolding errors, compared to the next best standalone scaffolder (1.5× SSPACE for *M. undulatus*, 3.5× SSPACE for *C. sinensis* and 1.4× SOPRA for *P. stipitis*).

To compare OPERA-LG with a wider range of scaffolding programs, we evaluated it on three additional datasets from a recent benchmarking study [11]. Contigs used in this study were corrected for assembly errors providing a more idealized setting for comparing scaffolders. Overall, OPERA-LG was consistently one of the top 3 methods in terms of corrected N50 across datasets (**Figure 5b, c, d**). No other method exhibited similarly consistent performance. For the two smaller genomes (*S. aureus* and *P. falciparum*), the datasets contained a single mate-pair library (2.8 kbp and 3.6 kbp respectively) and OPERA-LG had slightly lower corrected N50 than the best performing methods (1.1-1.2× improvement for MIP and ABYSS, respectively). Two mate-pair libraries (3 kbp and 35 kbp) were available for the third experiment (*H. sapiens* chromosome 14) and OPERA-LG provided significantly improved corrected N50 compared to other methods in this setting (>2×; **Figure 5d**). In terms of the number of incorrect joins (used to measure scaffolding error), no single method performed consistently well. Using an alternate metric [11] that takes the number of correct scaffold joins as a proxy for assembly contiguity, and computes a weighted sum with the number of incorrect joins to obtain a “normalized score”, SOPRA was observed to be the best method in 2 out of 3 datasets (**Supplementary Figure 4**). Note that the number of correct joins does not correlate well with assembly contiguity (e.g. OPERA-LG and SOPRA have a similar number of correct joins on the *H. sapiens* chromosome 14 dataset but OPERA-LG provides corrected N50 that is 5× that of SOPRA; **Supplementary Figure 4e** and **Figure 5d**). Thus the normalized score from Hunt *et al*. [11] provides an alternate measure of scaffold quality compared to widely used measures of assembly contiguity (a primary goal for genome assembly) such as corrected N50.

**Figure 5:**
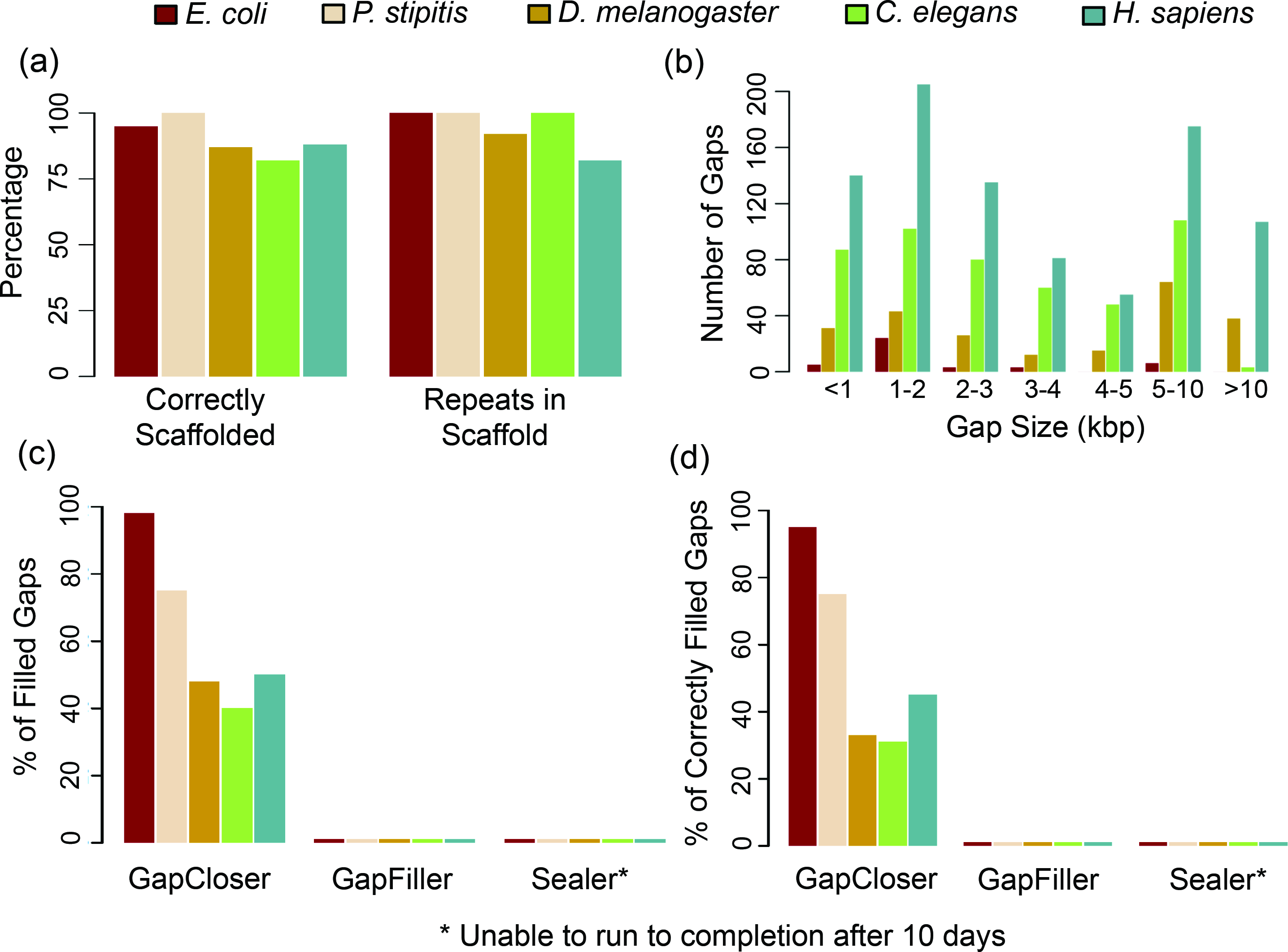
**Scaffolding of repeat sequences with OPERA-LG**. (a) Figures showing the correctness and completeness (“Repeats in Scaffold”) of scaffolding with repeat contigs in OPERA-LG. A set of contigs is considered correctly placed in a gap if the contigs belong to that gap and if they are in the right order and orientation. For completeness, all repeat contigs (longer than 500bp) in valid gaps (gaps where the adjacent contigs are in the right order and orientation) and with edges in the scaffold graph were considered. (b) Length distribution of gaps with contigs placed by OPERA-LG in various genomes. (c)-(d) Results from evaluating whether various gap-filling methods can substitute for OPERA-LG: (c) Percentage of gaps filled and (d) Percentage of correctly filled gaps, by different gap-filling methods in scaffolds gaps where OPERA-LG correctly placed a repeat contig. SEALER was unable to run to completion after 10 days.

Another aspect to note here is the incorporation of “skipped tags” (contigs omitted from a scaffold) as a component in the normalized score from Hunt *et al*. [11]. As the position of small contigs (default < 500bp) in a scaffold is frequently under-constrained, OPERA-LG omits them to avoid scaffold errors (**Supplementary Figure 4**), while still scaffolding most of the sequence (>94% of the genome for datasets from Hunt *et al*.). More importantly, the exclusion of very small contigs does not diminish the ability to correctly recover complete genes from OPERA-LG assemblies compared to the best assemblies based on the normalized score (**Supplementary Figure 5**). Paradoxically, the “normalized score” for OPERA-LG can improve using the complete set of contigs, even when corrected N50 and the number of incorrect links becomes worse (*P. falciparum* dataset; **Supplementary Figure 4d**; assembly size increase of 3%).

## Application I: Scaffolding of Repeat Sequences

Assembly of repeat sequences is typically the most error-prone stage of many assembly pipelines as was observed in the results for the GAGE [8] and Assemblathon [9] competitions (e.g. the gap-filling stage in SOAPdenovo). They are often handled in a post-scaffolding, gap-filling stage. Algorithmic extensions in OPERA-LG allow it to simultaneously scaffold unique and non-unique regions of the genome and thus more appropriately scaffold repeat-rich genomes – a feature that is not available in other scaffolders which filter repeats during their preprocessing stage. Evaluation of scaffold correctness for OPERA-LG in the presence of repeat contigs showed that repeat contigs were placed in the correct order (as suggested in the reference genome; **Figure 5a**) in more than 80% of scaffold gaps (regions with no unique contigs). In addition, more than 90% of repeat contigs in gap regions were placed in OPERA-LG scaffolds (**Figure 5a**), highlighting the completeness of scaffolds despite the conservative placement of repeats in OPERA-LG (see **Methods**).

Repeat contigs scaffolded by OPERA-LG were found to be frequently part of large gaps in the scaffold (for *D. melanogaster*, about half of the gaps were longer than 5 kbp; **Figure 5b**). Large scaffold gaps present a challenging scenario for many gap-filling programs [32]. As no other scaffolder allows scaffolding of repeat sequences, we assessed OPERA-LG’s utility by testing the ability of three gap-filling programs (GapCloser: part of the SOAPdenovo package, GapFiller [33] and the recently published program Sealer [34]) to determine the sequence for scaffold gaps where repeats were placed by OPERA-LG. For *E. coli* and to a lesser extent the *P. stipitis* genome, GapCloser was able to provide the correct sequence for most of the scaffold gaps (>60%, **Figure 5c, d**). GapFiller did not correctly fill many gaps (1%) with repeats placed by OPERA-LG (it typically reported sequences for gaps <800bp long) and Sealer was not able to complete after 10 days on any of the 5 data sets. Sealer’s results were surprising but potentially due to the complexity of the assembly graph around repeats and the exhaustive search that it needs to identify the correct sequence. These results show that the placement of repeat contigs in large genomes by OPERA-LG frequently provides information that is not available from gap-filling programs (**Figure 5c, d**).

## Application II: Assembly Augmentation and Hybrid Assembly

With the increasing availability of new sequencing technologies and additional sequencing datasets at reduced costs, improvement of older draft genome assemblies in a systematic fashion is an area of increasing interest in the field. As the underlying algorithms in OPERA-LG are conservative and intended to minimize discordance with data, the use of OPERA-LG as an *assembly augmentation* tool is attractive and was explored further in two recent genome assembly projects. In the first project, for the assembly of a sweet orange genome (*C. sinensis*) [31], all available Illumina sequencing datasets were used with SOAPdenovo to generate a preliminary draft assembly (N50 of ∼430 kbp). This assembly was then augmented using OPERA-LG, with SOAPdenovo scaffolds as starting sequences and by reusing the larger mate-pair libraries (10 kbp and 20 kbp). The final N50 obtained by this process was four times larger (∼1.6 Mbp; **Figure 4e**) and the assembly was validated to be of high-quality using BAC-end sequences, genetic linkage maps and cytogenetic analysis [31]. In another project, an existing reference genome assembly for a Chinese hamster ovary cell line (CHO-K1) [35] was augmented with newly generated Illumina sequencing datasets (300 bp library at 61X and 10 kbp library at 16X base pair coverage). The resulting assembly boosted N50 6-fold (1.2 Mbp to 6.9 Mbp; **Figure 4e**) with >90% of the genome assembled into ∼500 scaffolds. The scaffolds were also used to close gaps *in silico*, filling >130,000 gaps and scaffold correctness was confirmed using alignments of the transcriptome to the assembly (Yusufi *et al*., in preparation).

As an example of a novel direction for hybrid assembly with OPERA-LG (in addition, to its ability to mix paired-read libraries and assemblies from different technologies), scaffolding with long reads from third-generation sequencing technologies was explored (by generating mate-pair libraries *in silico*; see Methods). OPERA-LG was evaluated on reads generated on the Pacific Biosciences (PacBio) and Oxford Nanopore (ONT, https://www.nanoporetech.com/) platforms and compared against two recently described long read scaffolders: SSPACE-LongRead (SSPACE-LR) [28] and LINKS [29]. All methods were tested on two synthetic (*D. melanogaster* and *C. elegans* genomes) and three real datasets (*S. cerevisiae, D. melanogaster* and *M. undulatus* genomes; Supplementary Table 6). For synthetic data, OPERA-LG and SSPACE-LR had similar corrected N50s, though OPERA-LG had significantly fewer errors (3-4×; Supplementary Table 6). While LINKS reported no errors on these datasets, its corrected N50 was 4-10× smaller than that for OPERA-LG (Supplementary Table 6). On the real ONT dataset, OPERA-LG improved corrected N50 of the original *S. cerevisiae* assembly by 3 fold (from 49 kbp to 158 kbp) and provided better corrected N50 compared to other approaches (1.15×). ONT reads were converted into scaffold links using a PacBio specific mapper in SSPACE-LR, and using a nanopore-specific mapper could further improve results [36]. On datasets with real PacBio reads, OPERA-LG improved corrected N50 for the *D. melanogaster* genome from 58 kbp to 1,290 kbp, a 7× improvement over LINKS, and a 2.4× improvement over SSPACE-LR with slightly fewer errors (498 vs 568; **Supplementary Table 6**). For the larger *M. undulatus* genome, OPERA-LG was the only scalable method (**Supplementary Table 6**). Despite the low coverage from long reads (∼3X for reads longer than 2 kbp) for this dataset, OPERA-LG improved corrected N50 by 4-9× depending on how scaffold edges were constructed (**Supplementary Table 6**). Runtime for OPERA-LG and LINKS was at most 6 hours and 1 hour respectively, while SSPACE-LR took >5 days for genomes longer than 100Mbp. OPERA-LG’s results provide a proof-of-concept for this application, and refinements in mapping and scaffold edge construction could further improve results for low coverage datasets.

## Discussion

For many bioinformatics problems it is either hard to formalize a clear objective for algorithm design, or the formalized objective is computationally intractable. Benchmarking of a novel algorithm is thus the norm for demonstrating the algorithm’s utility and advance over state-of-the-art. Due to resource limitations, however, benchmarks will always be limited in nature. Moreover, consensus on a standardized and comprehensive benchmark can be elusive in many areas, though recent efforts have provided valuable resources for benchmarking scaffolders and assemblers. Where feasible, algorithms that have a clear optimization criterion provide users reason to believe that the algorithm will work on a new dataset. This is the motivation behind OPERA-LG, where the algorithm is guaranteed to produce a minimal-repeat scaffold that globally minimizes the number of discordant scaffold edges. This is by no means the only optimization criterion though and algorithms such as SOPRA optimize very similar criteria in a probabilistic setting.

Results for OPERA-LG on synthetic and real datasets indicate that it is well suited for scaffolding datasets where multiple mate-pair libraries or large-fragment libraries are available. In such settings, OPERA-LG reported the best corrected N50s compared to other scaffolders and assemblers. With fewer libraries or smaller fragment libraries, the improvements seen were more limited, though OPERA-LG provided consistently good results. While corrected N50 is widely used to evaluate scaffolds and assemblies [8–10] other metrics may prefer alternate scaffolders (e.g. SOPRA using “normalized score” [11]). Deciding which method to use in an assembly project would thus likely depend on the specific requirements of the project.

OPERA-LG relies on the presence of sufficient mate-pair information to constrain the scaffold graph and report an optimal scaffold. When this is feasible, it provides scaffolds with fewer errors compared to other methods, particularly translocation and inversion errors. However, availability of only a single mate-pair library and limited sequence coverage can under-constrain the scaffolding problem in some regions, especially in the presence of short contigs. There can be multiple optimal scaffolds in such situations, and thus lead to mis-assemblies with OPERA-LG. OPERA-LG tries to mitigate this issue by selecting a solution based on approximate distance information from scaffold edges (see **Methods**). Additional strategies, such as post-processing to detect regions of the scaffold that are under-constrained and local analysis to correct misassemblies, or the assignment of a confidence score to different regions of the scaffold, deserve further consideration.

A common concern and criticism for exact algorithms for assembly and scaffolding has been that they are slow, not practical for large datasets and cannot handle complicated scenarios (e.g. for scaffolding repeats). An important contribution of this work is to show that these challenges are surmountable in a specific deterministic scaffold assembly framework with appropriate algorithm design and engineering. Our results show that large and repeat-rich genomes are amenable to assembly using exact algorithms and that the efficiency of OPERA-LG is comparable to other scaffolding methods. Methods such as SOPRA and OPERA-LG could thus allow users to obtain consistently high quality scaffolds, depending on the scaffold quality metric of choice.

## Conclusions

As characteristics of sequencing technologies continue to improve, particularly in read length, assembly quality is also expected to benefit [37, 38]. Recent improvements in protocols for the generation of large mate libraries (e.g. 40 kbp libraries from the NxSeq technology: http://www.lucigen.com/NxSeq-DNA-Sample-Prep-Kits/), further emphasize the need for effective scaffolding methods that can use this information. However, genome assembly will continue to be a trial-and-error process until assembly and scaffolding algorithms fully exploit the power of the data. Assembly errors continue to plague downstream biological sequence analysis, affecting almost every aspect of modern bioinformatics, and a full assessment of their impact has yet to be performed [6]. Our analysis suggests that commonly used assembly tools such as SOAPdenovo and SSPACE can introduce hundreds to thousands of errors during scaffolding. As these programs have been used to construct hundreds of draft eukaryotic genomes, they may benefit from re-assembly and assembly augmentation using newer assembly tools. As we move to an era where assembled sequences potentially guide medically relevant decisions, tolerance for assembly errors is likely to be even more limited. Improved assembly tools are thus needed, such that, fully-phased and complete, quality-guaranteed assemblies are the norm in the future.

## Methods

### Constructing the scaffold graph in OPERA-LG

In order to simultaneously use data from multiple “jumping” libraries for scaffolding, OPERA-LG uses a three-staged process to combine library information to construct a scaffold graph (with contigs as nodes and scaffold edges, that define the constraints on order, orientation and distance between adjacent contigs, linking them). Firstly, instead of relying on user input for library properties such as mean and standard deviation of insert sizes, as well as read orientation, by default, OPERA-LG directly estimates them from read mappings on large sequences (>2× the insert size, as estimated from a set of 1,000 read-pairs). The read orientation of a library is set by default to the majority orientation of all read-pairs mapped to the same contig and these are then used to calculate the mean insert length and standard deviation. To avoid biases due to outliers (from mis-assemblies, mis-mapping or sequencing errors), read-pairs with distance lesser than *Q*_1_ – 3 x *IQR* or greater than *Q*_3_ + 3 x *IQR* were ignored, where *Q*_1_ and *Q*_3_ are the first and third quartiles respectively of distances between read-pairs on a contig and *IQR* = *Q*_3_ – *Q*_1_.

#### Refined Edge Length Estimation

Secondly, OPERA-LG combines information from read-pairs in a single library to get a library specific estimate of edge length for each scaffold edge. Let the gap length between two contigs be *g*, and *C* be the sum of the contigs length and *g*. To account for the fact that the observed read-pairs are from a truncated distribution [30] (in the range *[g, C]* as shown in **Supplementary Figure 6**), a reverse lookup table is used to estimate *g* (gap size) from the observed mean of *Ŝ* (defined as the length of contig sequences that overlap with the insert region of the paired-reads; see **Supplementary Figure 6**). As noted in Sahlin *et al*. [30], this is an important step to avoid biases in edge length estimation that could lead to incorrect ordering of contigs and we propose a novel approach for bias correction. Specifically, given *g* and the overall distribution of insert lengths *I* (as determined by the read mapping on large sequences), we compute *E*(*Ŝ*) as *E*(*I*_[*g,c*]_) – *g* (where *I*_[*g,c*]_ is the random variable for the truncated distribution indicated in **Supplementary Figure 6b**) for every value of *g* in the range [0, *L*] (where *L* is an upper bound of the library insert size, by default *μ* + 6σ). To reduce runtime, OPERA-LG pre-computes such a lookup table for *l* (= sum of contig lengths) in the range [*g, Q*_3_ + 3 x *IQR*] at 500bp intervals and returns the value of *g* that corresponds to the value of *E*(*Ŝ*) closest to the observed mean of *Ŝ* (using the appropriate lookup table for the current value of *l*). Note that the pre-computation for a lookup table is quite efficient as it can be done in linear time (as a function of the library size) using cumulative sums.

#### Multi-library Scaffolding

Thirdly, scaffold edges obtained from different libraries are combined to form a unified scaffold graph in OPERA-LG. To do this, OPERA-LG uses the mean (as obtained above) and standard-deviation [21] estimates for all edges connecting a pair of contigs (in the same orientation) to cluster and merge edges, starting at each stage with the edge with the largest standard deviation and identifying other edges whose means are within *k* (= 6 by default) standard deviations of this edge to merge into a single edge. In the case where more than one edge remains at the end of the process, OPERA-LG uses the edge supported by the most paired-reads and discards other edges (for assembly of polyploid sequences all edges will be discarded and for repeat contigs all edges will be retained).

Since libraries with very different insert sizes (say 200 bp vs 40 kbp) provide largely orthogonal information for scaffolding, it is possible to consider them in groups of similar insert sizes. OPERA-LG therefore allows the user to combine libraries in a *staged* fashion (< 1kbp, 1-10kbp and > 10kbp, by default) for greater runtime efficiency.

## Scaffolding of Unique Sequences

OPERA-LG is based on Opera [13], which is restricted to the scaffolding of unique sequences. In this section, we briefly describe the main concepts that are common to Opera and OPERA-LG. We begin with a few definitions. We define a *scaffold* (without repeats) as being given by a signed permutation (where the sign denotes orientation) of unique contigs as well as a list of gap sizes between adjacent contigs. A scaffold edge *e* = < *c*_1_,*c*_2_ > is a concordant edge if *c*_1_ and *c*_2_ in the scaffold can satisfy the distance and orientation constraints imposed by *e* and otherwise *e* is marked discordant. Given a scaffold graph *G* = (*V, E*), a partial scaffold *S*’ is a scaffold on a subset of the contigs and the dangling set, *D*(*S*’), is composed of edges from *S*’ to *V* – *S*’. The active region *A*(*S*’), is given by the shortest suffix of *S*’ such that all dangling set edges are adjacent to a contig in it. A partial scaffold *S*’ is considered *valid* if all edges in the induced subgraph are concordant.

**Definition 1 (Scaffolding Problem Without Repeats):** Given a scaffold graph *G*, find a scaffold *S* of the contigs that maximizes the number of concordant edges.

This problem is analogous to that in Huson *et al*. [21] (where the problem is defined in terms of individual paired-reads), and therefore the proof there can be adapted in a straightforward way to show that the decision version of the scaffolding problem is NP-complete.

We first consider the special case where the optimal scaffold in a scaffold graph has no discordant edges. Avoiding a naïve and intractable search over all possible signed permutations, we can instead limit the search to an equivalence class of partial scaffolds as shown in the following lemma from Gao *et al*. [13].

**Lemma 1**. If 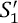 and 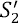 are two valid partial scaffolds where 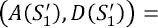 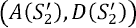 then (1) 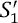 and 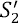 contain the same set of contigs; and (2) both or neither of them can be extended to a valid complete scaffold.

Hence, the scaffolding algorithm of Opera without any discordant edges (as defined in Gao *et al*. [13]) requires *O*(|*E*||*V*|^*w*^) time, where *w* as an upper-bound on the number of contigs that can be spanned by a paired-read.

Treating the total number of discordant edges *p* in the scaffold graph as a constant, we can extend the previous algorithm and still maintain a runtime polynomial in the size of the graph. To do this, we extend the notion of equivalence class by keeping track of discordant edges from the partial scaffold (denoted by *X*(*S*’) for a partial scaffold *S*’). We also redefine the notion of dangling set to only contain concordant edges. As the scaffold is only extended to the right of the active region, given a certain active region it is possible to collect all edges connecting to its right side. Thus, the dangling set is obtained by subtracting the discordant edge set from all edges adjacent to the active region. The following lemma is then a straightforward extension of Lemma 1:

**Lemma 2**. If 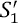 and 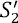 are two partial scaffolds with less than *p* discordant edges and 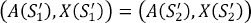 then (1) 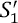 and 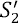 contain the same set of contigs; and (2) both or neither of them can be extended to a valid complete scaffold.

The extended scaffolding problem with *p* discordant edges can be solved in *O*(|*E*|^*p*+1^|*V*K|^*w*^) time as shown in Gao *et al*. [13].

**Supplementary Figure 1** shows the core common features of Opera and OPERA-LG. The memoized-search for an optimal scaffold based on **Lemma 2** treats all contigs as possible starting points (**Supplementary Figure 1a**). Such a search can be localized using a graph contraction approach to reduce runtime. The basic idea here is that a successful contig assembly would frequently produce contigs that are much longer than the paired-read library threshold(s). If we label such contigs as *border contigs* and note the fact that a valid scaffold will not have concordant library edges spanning a border contig, then for a scaffold graph *G* = (*V,E*), we can define *G*_0_ = (*V*_0_, *E*_0_) as a fenced subgraph if edges in *E* from *V* — *V*_0_ to *V*_0_ are always adjacent to a border contig. The search can then be localized in this fenced subgraph without losing the guarantee of a globally optimal scaffold [13] (**Supplementary Figure 1b**). After the order and the orientation of contigs in a scaffold has been computed, the size of intervening gaps is determined based on constraints imposed by the paired-reads. As scaffold edges can span multiple gaps, imposing competing constraints on their sizes, a maximum likelihood approach was adopted to compute gap sizes optimizing a clear likelihood function while taking all scaffold edges into account [13] (**Supplementary Figure 1c**).

## Scaffolding of Repeat Sequences

OPERA-LG relies on the read coverage of contigs to identify potentially repetitive contigs in the assembly (as well as polyploid sequences, as detailed in **Supplementary Note 1**). More sophisticated approaches that use the graph structure to do this (as is done in the Celera Assembler) could potentially yield more accurate and sensitive results (especially in low coverage settings) and will be explored in future versions of OPERA-LG. Since coverage estimates may not always be provided directly by the assembler (e.g. ALLPATHS [26]) or can be inaccurate (e.g. SOAPdenovo k-mer coverage values are bounded at 63), OPERA-LG computes these directly from user-provided read mappings. For haploid genomes, OPERA-LG identifies sequences with coverage less than a multiple (1.5 by default^1^) of the genomic average as *unique*. Here, we describe how the search procedure in Gao *et al*. [13] can be extended to simultaneously scaffold repeat sequences and unique sequences. There are two assumptions for handling repeats in our algorithm: (1) Repeats cannot be used to extend scaffolds – this is to avoid ambiguous extensions to the scaffold and (2) The concordance of edges between repeats is ignored as these cannot be verified without assuming that all repeat instances have been scaffolded.

### Definitions

To incorporate repeat contigs, we slightly extend definitions presented in the previous section: A *scaffold* is given by a signed permutation of the contigs (in which repeat contigs are allowed to occur multiple times) as well as a list of gap sizes between adjacent contigs. An edge *e* =< *c*_1_ *c*_2_> is a *concordant* edge if any pair of instances of *c*_2_ and *c*_2_ in the scaffold can satisfy the distance and orientation constraints imposed by *e* and otherwise *e* is *discordant*. For a scaffold graph *G* = (*V, E*) and a given partial scaffold *S*’, the *dangling edge set D* (*S*’) is the set of edges from unique contigs in *S*’ to all contigs in *V* — *S*’. When the distance between a repeat *r* and the tail of *S*’ (measured as sum of contig lengths in the partial scaffold) is larger than the upper-bound of the paired-read library, *r* is said to be *confirmed* (in the sense that it will not be removed from the partial scaffold constructed by OPERA-LG). Then we define the *active region A*(*S’*) as the shortest suffix of *S’* (including all unconfirmed repeats) such that all concordant dangling edges are adjacent to a contig in *A*(*S’*). Also, we use the notation *u(G)* and *u(S)* to refer to the subgraph of *G* and the subset of *S* composed of unique contigs only.

To constrain the number of occurrences of repeat contigs in scaffolds, we use a simple parsimony criterion to redefine our notion of an optimal scaffold:

**Definition 2 (Minimal-repeat Optimal Scaffold)**. *Given a scaffold graph G* = (*V,E*) *and a scaffold S that minimizes the number of discordant edges in the graph, S is considered a “minimal-repeat optimal scaffold”, if removing any occurrence of a repeat from S will increase the number of discordant edges*.

Correspondingly, we have the following updated formulation of the Scaffolding Problem with Repeats:

**Definition 3 (Scaffolding Problem with Repeats)**. *Given a scaffold graph G, find a minimal-repeat optimal scaffold S of the contigs*.

Note that the criterion for including repeats is inherently conservative and would, for example, favor the placement of a single copy of a tandem repeat, where feasible. Suitable post-processing scripts would therefore be needed to estimate and expand-out copies of a tandem repeat region. We next describe how OPERA-LG’s search procedure is designed to guarantee a scaffold that minimizes discordance with paired-read derived scaffold edges while parsimoniously including repeat contigs in the scaffold.

### Construction of a minimal-repeat optimal scaffold

As is the case for unique sequences, OPERA-LG extends partial scaffolds only to the right of the active region, and all contigs are tried as potential starting points. Also, as before, the search in OPERA-LG is limited to an updated equivalence class of partial scaffolds as follows:

**Lemma 3**. Given a scaffold graph G and two valid partial scaffolds *S’*_1_ and *S’*_2_ with *k*_1_ and *k*_2_ discordant edges respectively, if (*A*(*S’*_1_), *D*(*S’*_1_),*k*_1_) = (*A*(*S’*_2_), *D*(*S’*_2_),*k*_2_) then (1) *S’*_1_ and *S’*_2_ contain the same set of unique contigs; and (2) both or neither of them can be extended to a solution with equal or less than *k* discordant edges (∀*k* ∈ ℕ).

*Proof*. For (1), since the dangling edge set defines a cut in *u(G)*, *D*(*S’*_1_) = *D*(*S’*_2_) define the same cut and since (*S’*_1_) = *A*(*S’*_2_), *S’*_1_ and *S’*_2_ must be on the same side of this cut and thus contain the same set of unique contigs.

For (2), let *S"* be any scaffold extension of *S’*_1_ such that *S’*_1_ has ≤ *k* discordant edges. Then *S’_2_ *S"* would also be a valid scaffold as *u*(*S"*) = *u*(*V* – *S’*_1_) = *u*(*V* — *S’*_2_). Also, since the active regions are identical, any newly discordant edge in *S’*_1_ *S"* (i.e. not discordant in *S’*_1_) that is adjacent to a contig in *S"* will also be newly discordant in *S’*_2_ *S"* (i.e. not discordant in *S’*_2_)* and *vice versa*. The corresponding number of discordant edges in *S’*_2_ *S"* is ≤ *k*_2_ + (*k* — *k*_1_) = *k*.

During the memorized search in OPERA-LG (see **Supplementary Figure 1a**), a minimal-repeat optimal scaffold is obtained based on the following definition and lemma:

**Definition 4 (Essential Repeat Instance)**. A repeat instance *r* in a partial scaffold is considered essential if no extension of the partial scaffold can be optimal if *r* is removed.

**Lemma 4**. (a) A repeat instance *r* in a partial scaffold *S* is essential *iff* (b) removing *r* increases the number of discordant edges when *r* is being confirmed (i.e. in the process of being tested to see if it should be marked confirmed).

*Proof*. Let the partial scaffold *S’* be obtained by removing *r* from *S*. (1) To prove (*a*) ⇒ (*b*), for the sake of contradiction, let a repeat instance *r* in a partial scaffold *S* be essential s.t. removing *r* does not increase the number of discordant edges when *r* is being confirmed. Then, if there exists an optimal extension *T* of *S* with *p* discordant edges, the scaffold *S’T* also has at most *p* discordant edges (as the status of edges connecting to *T* cannot change from *ST* to *S’T*, from the definition of a confirmed repeat) and is therefore also optimal. Hence *r* is not essential, giving a contradiction. (2) To prove *(b)* ⇒ *(a)*, suppose removing a repeat instance *r* increases the number of discordant edges when *r* is being confirmed. Then for any extension *T* of *S’* s.t. *S’T* has *p* discordant edges, the scaffold *ST* will have less than *p* discordant edges (as before, the status of edges connecting to *T* cannot change) and hence, *S’T* can never be optimal. By Definition 4, therefore, *r* is essential in *S*.

Note that by definition, non-essential repeat instances should not be included in a minimal-repeat optimal scaffold. Correspondingly, based on Lemma 4, the algorithm ScaffoldWithRepeat presented in **Supplementary Figure 7** is guaranteed to report a minimal-repeat optimal scaffold.

The runtime complexity for ScaffoldWithRepeat is established in the following theorem.

**Theorem 1**. Consider a scaffold graph *G* = (*V,E*). Let *p* be the maximum allowed number of discordant edges and *w* be the maximum number of contigs in the active region. The algorithm ScaffoldWithRepeat runs in *O*(*p*|*V*|*^w^*|*E*|^*p*+1^) time.

*Proof*. It is straightforward to show that the set of possible active regions is *O*((2|*V*|)^*w*^) and there are at most *O*(2^*w*^2^^) possible sets of concordant dangling edges for any given active region. In addition, there are at most *O*(|*E*|^*p*^) possible sets of discordant dangling edges. So, there are at most *O*(|*E*;|^*p*^2^*w*^2^^) = *O*(|*E*;|^*p*^) sets of dangling edges. The number of equivalence classes is therefore bounded by *O*((2|*V*|)^*w*^|*E*|^*p*^*p*) = *O*(*p*|*V*|*^w^*|*E*|^*p*^). For each equivalence class, confirming repeats, updating the active region, the dangling set and the number of discordant edges in steps 6-11 takes *O*(|*E*|) time.

As is the case for scaffolding unique sequences, we use the concept of fenced subgraphs and graph contraction to improve the runtime of OPERA-LG in practice, without affecting its guarantee of finding an optimal scaffold. Note that since repeats are not used for extending scaffolds, we can construct fenced subgraphs on the unique subgraph *u(G)* in the same manner as for unique sequences. Then, the *fenced subgraph with repeats* can be obtained by adding back repeat contigs with edges to unique contigs in a subgraph. The results on unique sequences can then be extended trivially to show that finding minimal-repeat optimal scaffolds on fenced subgraphs with repeats will lead to a global optimum on the entire graph. For polyploid genomes, OPERA-LG uses an intuitive (but untested) approach to accommodate for repeats, and thus avoid the scaffolding errors that they can induce (see **Supplementary Note 1**).

## Scalability and Optimized Data Structures

Due to the computational intensity of the search procedure employed in OPERA-LG [13], engineering issues such as code optimizations and data structures play an important role in enabling its application to large genomes. To allow for greater control over runtime and memory optimizations, OPERA-LG is implemented in C++ with custom data structures designed to allow for a smaller memory and runtime footprint by exploiting the depth-first structure of the search [13]. Specifically, as shown in **Supplementary Figure 1a**, the search procedure in OPERA-LG needs to keep track of a partial solution *S* during its search by recording the corresponding active region *A(S)* and a set of discordant edges *X(S)*. Since partial scaffolds that are related in the search tree (say *S*_1_ and *S*_2_) can overlap in their respective active regions and discordant edge sets, OPERA-LG avoids duplication of this information, by only recording the difference in these sets (i.e. *A*(*S*_1_) △ *A*(*S*_2_) and *X*(*S*_1_) △ *A*(*S*_2_)). In addition, as shown in **Supplementary Figure 1a**, the search avoids unnecessary computation by keeping track of partial scaffolds that have already been explored (memoization). While this can be done in a straightforward way using a hashtable, memory requirements for this approach can be significant. In OPERA-LG we implemented a *prefix tree* data structure to store active regions (these are lists by definition) and corresponding discordant edges (an arbitrary order was imposed to convert these sets into lists) as shown in **Supplementary Figure 8**. This allowed for a significant reduction in the memory footprint of OPERA-LG (**Table 1**) while allowing for lookups in time proportional to the size of the active region and the discordant edge set.

While the runtime requirements for OPERA-LG were typically found to be modest and in particular aided by the graph contraction step shown in **Supplementary Figure 1b**, the search time for some sub-graphs can be significantly longer than average. Correspondingly, OPERA-LG allows the user to bound the number of partial scaffolds enumerated on any one sub-graph (default value is 1 million), switching to solving the problem with an increased edge size threshold (default step size of 1) when the maximum is reached. This hybrid-exact option in OPERA-LG (default setting) allows for the user to benefit from an exact algorithm for most of the assembly (>90% of the genome in all datasets tested here) while relying on a reasonable heuristic (edges with few supporting reads are less reliable) when an exact approach is not directly feasible. Though the rate of incorrect scaffold joins for such sub-graphs was found to be comparable to the rest of the graphs (0.5% in both cases), OPERA-LG conservatively flags such scaffolds to the user for further investigation.

Finally, to improve correctness on large genomes with shallow read coverage, the search procedure in OPERA-LG explores contig extensions in increasing order of their estimated distance from the end of a partial scaffold (line 4 in **Supplementary Figure 7**). To estimate these distances, it employs a breadth-first-search (visiting each scaffold edge only once) and uses a weighted mean (down-weighting by the number of edges) for a contig that can be reached by more than one path. This ordering of contig extensions does not impact the performance guarantees in OPERA-LG (i.e. its ability to find an optimal scaffold) but typically generates more accurate scaffolds in regions with ambiguous extensions.

## Hybrid Assembly and Scaffolding with Long Reads

Similar to other scaffolders that are not restricted to a specific mapper or assembler [14], OPERA-LG allows users to combine contig assemblies and paired reads from different sequencing technologies. This feature of OPERA-LG has been exploited in several projects including the assembly of CHO cell lines using SOLiD mate-pair and Illumina paired-end datasets (Yusufi *et al*., manuscript in preparation). In addition, the availability of ‘third-generation’ sequencing technologies that directly produce longer reads (e.g. median lengths in the range 2-8 kbp from PacBio Systems [37]) has provided another avenue for significantly improving sequence contiguity during genome assembly. For example, in the case of the *M. undulatus* assembly, the authors reported significant improvement in contig N50s (30-fold to 100 kbp) by including PacBio reads [38]. The use of PacBio reads to aid scaffolding has also been demonstrated by custom methods designed for this task (SSPACE-LongRead [28] and LINKS [29]). As a proof of concept that OPERA-LG can adapt to this task with minimal modification, we tested it with contig links (i.e. synthetic mate-pairs from long reads) inferred by SSPACE-LongRead as input (see **Supplementary Note 2** for an alternative approach). For each contig link we fixed the standard deviation in distance as 10% of the distance estimated by SSPACE-LongRead (to account for indel errors in PacBio and ONT reads), grouped links into synthetic mate-pairs libraries (with estimated distances in the range [0-300], [300-1,000], [1,000-2,000], [2,000-5,000], [5,000-15,000] and [15,000-40,000]), and provided the synthetic libraries as input to OPERA-LG to construct scaffold edges and link contigs together into scaffolds.

## Evaluation on Synthetic Datasets

All paired-end and mate-pair read libraries for synthetic datasets were generated using Metasim with default Illumina sequencing settings [39]. In addition to the three small genomes (*E. coli, S. cerevisiae* and *D. melanogaster* chromosome X) analyzed in Gao *et al*. [13], we generated libraries and benchmarked on three larger genomes as well i.e. *D. melanogaster, C. elegans* and *H. sapiens* (reference genomes from NCBI with details in **Table 2**). PacBio reads were generated using PBSIM v1.03 [40] (using --data-type CLR, i.e. long reads with high error rate) to mimic a library of 3 kbp mean read length (maximum read length of 25 kbp) and give ∼7X coverage of the *D. melanogaster* and *C. elegans* genomes.

For the large synthetic datasets, the GAGE pipeline was used to evaluate the final assemblies [8] for contig errors and scaffold errors (indels longer than 5bp, inversions, relocations and translocations). As the terminology can be confusing, it is important to note here that scaffold indel errors in GAGE are defined as regions between two contiguously scaffolded contigs A and C, where a contig B larger than 200bp could be placed, and where the gap estimation between A and C is not within 1kbp of the size of B. Corrected assemblies were produced by splitting the final assemblies produced by all programs at contig and scaffold errors (for evaluating the overall assembly) or only scaffold errors (for scaffold-level evaluation) to compute the *corrected N50* (N50 is defined as the fragment length such that >50% of the genome is in fragments of equal or longer length). For handling scaffolds with repeats, read mappings produced by the GAGE pipeline were further analyzed. A mapping position was considered correct if both coverage and identity was found to be greater than 90%. For each repeat, all correct positions were considered for correctness and completeness analysis. To evaluate the correctness of gap-filled sequences, we aligned them to the corresponding reference sequence (using MAFFT with default parameters [41]). A gap was considered correctly filled if similarity was >95% and length difference <5%.

## Benchmarking on Sequenced Genomes

Publicly available data from seven sequenced and published genomes were further used to benchmark scaffolders and assemblers in this study: a bacterial genome (*Staphylococcus aureus*, 2.8 Mbp) [8, 11], two yeast genomes (*Saccharomyces cerevisiae*, 12Mbp and *Pichia stipitis*, 15.0 Mbp) [42], a parasite genome (*Plasmodium falciparum*, 23 Mbp) [11], a mammalian chromosome (*Homo sapiens* chromosome 14, 100 Mbp) [11], a fruit genome (*Citrus sinensis*, 367 Mbp) [31] and a bird genome (*Melopsittacus undulatus*, 1.2 Gbp) [10].

Due to the availability of a high-quality reference for *P. stipitis* (NCBI accession number NZ_AAVQ01000000) and *S. cerevisiae* (http://www.yeastgenome.org/ strain S288C) assemblies were evaluated using the GAGE pipeline. For *S. aureus, P. falciparum* and *H. sapiens* chromosome 14, assemblies were evaluated using scripts provided by Hunt *et al*. [11], and corrected N50 values were computed by breaking assemblies at incorrect joins. For other genomes, the assemblies were evaluated for errors using REAPR [43] (used as part of the evaluation pipeline for Assemblathon2 [10]). For *C. sinensis* and *M. undulatus*, all reads from the 10 and 20 kbp mate-pair libraries, respectively, were used for REAPR evaluation (mapped using –i 15000). For *M. undulatus*, PacBio reads were mapped onto the assemblies to recognize contig errors, defined as regions not covered by *any* PacBio reads and having more than three split-mapped reads within 100 bp of the putative breakpoint. Reads were aligned to the assembly using Nucmer v3.23 (with –maxmatch), and the best mapping position for each read was selected using delta-filter v3.23 with default parameters. This approach resulted in only 1 false-positive break on a simulated dataset with 7X coverage of the SOAPdenovo contig assembly for *M. undulatus*. Corrected assemblies were produced for all datasets by splitting the final assemblies produced by all programs at contig and scaffold errors (for evaluating the overall assembly) or only scaffold errors (for scaffold-level evaluation) to compute the corrected N50.

## Parameter Settings

Parameter settings for the benchmarked scaffolders and assemblers were as follows – SSPACE (SSPACE-BASIC-2.0): default command line parameters, 6 * standard deviation / library mean was used as threshold for allowed error for each library; BESST (version 1.3.9): default parameters; SOPRA (version 1.4.6): run using a wrapper script from Hunt *et al*. [11]; SOAPdenovo2 (version 2.04): -K 41, -d 1, map length was fixed to half of the read length for each library; and ALLPATHS-LG (version 43019): PATCH_SCAFFOLDS=false, KPATCH=false (to skip the gap-filling stage). OPERA-LG was run using default parameters: contigs smaller than max(500, 2 × paired-end insert-size) bp were not scaffolded, contigs with coverage greater than 1.5 times the average coverage were treated as repeats and scaffold edges with <5 supporting reads were discarded. For runtime efficiency, all genomes >200 Mbp were scaffolded using OPERA-LG’s staged approach.

Contigs produced by SOAPdenovo (with -K 41 -d –D) were provided as input for SSPACE, BESST, SOPRA and OPERA-LG (except for datasets from Hunt *et al*. [11]). Read mappings for OPERA-LG and SSPACE was generated using BWA (0.7.10-r789) (samse -n 1) [44]. Read mapping for SOPRA and BESST were performed using the script provided in their respective packages and the same version of BWA. Assemblies from Assemblathon2 produced by Meraculous, BCM-HGSC and Newbler-454 were used for comparison as they were reported to be the best assemblies for *M. undulatus* based on several criteria [10]. For datasets from Hunt *et al*., read mappings for OPERA-LG and BESST were performed using BWA (0.7.10-r789) (samse -n 1) on the assemblies provided [11]. As multiple mappers were used to assess the performance of other methods, we conservatively used the best reported assembly.

For the gap-filling analysis, parameter settings were as follow: GapCloser (version 1.12): default parameters; GapFiller (version 1.10): -m 30 -o 2 -r 0.7 -n 10 -d 50 -t 10 -g 0 -i 1 (recommended parameters); and Sealer (version 1.9.0): -P 10 -k90 -k80 -k70 -k60 -k50 -k40 -k30 (recommended parameters) -j 20 (20 threads) -F 15000 (to allows filling of gaps up to 15 kbp long).

For scaffolding using PacBio and ONT reads we used SSPACE-LongRead (version 1.1) and LINKS (version 1.5.2) with default parameters. For low coverage datasets (simulated *D. melanogaster, C. elegans* and *M. undalatus*) OPERA-LG’s edge support threshold was set to 1, as chimera rates are expected to be low for contig links derived from PacBio reads. For ONT data, we excluded the template and complement reads of corresponding 2D reads.

## Availability of Data and Materials

Source code and executables for OPERA-LG are freely available from its SourceForge website: http://sourceforge.net/projects/operasf/. Simulated datasets and assemblies presented (except the ones that were downloaded from Assemblathon 2 and Hunt *et al*.) are available at the following ftp site: ftp://ftp2.gis.a-star.edu.sg/opera-lg/. Published sequencing datasets were accessed based on information from the following websites (1) *Melopsittacus undulatus*: Accession numbers from http://www.gigasciencejournal.com/content/supplementary/2047-217x-2-10-s5.xlsx were used to download data from the Sequence Read Archive (http://www.ncbi.nlm.nih.gov/sra), (2) *Pichia stipitis*: sequencing data was downloaded from ftp://ftp.jgi-psf.org/pub/JGI_data/meraculous/, (3) *Citrus sinensis*: sequencing data was downloaded from http://citrus.hzau.edu.cn/orange, (4) *Staphylococcus aureus*: data was downloaded from http://gage.cbcb.umd.edu/data/index.html, (5) *Plasmodium falciparum*: data for the runs ERR034295, ERR163027, ERR163028 and ERR163029 were downloaded from http://www.ebi.ac.uk/ena, (6) Human chromosome 14 data was downloaded from http://gage.cbcb.umd.edu/data/index.html, (7) *S. cerevisiae* W303 raw ONT sequences were from http://schatzlab.cshl.edu/data/nanocorr, (8) *D. melanogaster* PacBio sequences were from https://github.com/PacificBiosciences/DevNet/wiki/Drosophila-sequence-and-assembly.

## Authors’ Contributions

SG, DB and NN participated in designing the algorithm. SG implemented OPERA-LG. SG, DB and BC conducted the experiments and analysis with guidance from NN. SG, DB and NN drafted the manuscript. All authors read and approved the final manuscript.

## Additional Data Files

The following additional data are available with the online version of this paper: Additional data file 1 contains supplementary notes, figures and tables.

## Footnotes

1 Note that the A-statistic [27], with coverage modeled as a Poisson distribution, gives rise to a similar threshold (≈ 1.44).

## Acknowledgements

This work was supported by the IMaGIN platform (project No. 102 101 0025), through a grant from the Science and Engineering Research Council as well as funding to the Genome Institute of Singapore from the Agency for Science, Technology and Research (A*STAR), Singapore.

